# Integrated multi-omics data reveals the molecular subtypes and the contruction of novel signature to aid Immunotherapy

**DOI:** 10.1101/2023.03.09.531978

**Authors:** zhenqing li, Bin Li, Yuqi Meng, Kerong Zhai, Xuan Li

## Abstract

Malignant cancer exhibits a severe imbalance between the active of immune response and tolerance. And the different and connection between the establishment and deficiency of the tumor immune response is still unknown entirely. Here, we used ten clustering algorithms based on five immune cells infiltration matrix to identified two specific molecular subtypes and determine its potential clinical outcomes and immune response effect. The cluster A subtype can be called the cold inflammatory subtype, which revealed the worst prognosis. And the cluster B can be identified as hot inflammatory subtype with high activated immune-related pathways. Then we used the WGCNA algorithms and limma to confirm the driven genes between the cluster A and B. 67 genes were identified the cluster A signature genes, and based on that we used the ssgsea algorithm to quantize the two subtypes. We confirmed the high score group had high activated level of immune activity pathways and better survival rate. The tumorigenesis-associated pathways were analysed in low score group, indicating active cell proliferation may be associated with poor prognosis. An in-silico drug screen predicted the erlotinib can be an effective compound in low score group based on the lung cancer cell line. Our research confirmed that the score can worked as an effective immunotherapy index, which provide a promising therapeutic strategies on immune therapeutics for lung adenocarcinoma.

## Introduction

Lung cancer is the main reason of cancer-related death around the world with poor survival and highly malignant(1). And as the relevant statistics, there are 2.2 million new cancer patients was diagnosed and 1.8 million deaths in the worldwide (2). Lung cancer can be divided into two histological subtypes, including non-small cell lung cancer (NSCLC, which accounts for 85% of lung cancer) and small cell lung cancer (SCLC, which accounts for 15% of lung cancer). The NSCLC had three main histological types, including large-cell lung, lung adenocarcinoma (LUAD) and lung squamous-cell carcinoma (LUSC) (3). It is more and more clear that the NSCLC does not represent a single disease entity, and present a diverse set of molecular-driven tumors, which changed the landscape of NSCLC treatment for personalized treatment based on molecular changes. Despite these therapeutic advances, NSCLC is still can take place metastatic especially in the absence of an EGFR mutation or ALK translocation, which with a disappointing median overall survival in one year (22). The SCLC was characterized by rapid growth and a tendency toward early metastasis with aggressive clinical course (23). Despite highly sensitive to chemotherapy, most patients with SCLC would easily relapse and had little long-term survival rate(23).

In the recently years, the emergence of immunotherapiers brings immense efficiency in promoting the survival rate for patients (15). And the immunotherapy in the treatment of melanoma and renal cell carcinoma had been proved that it can improve effective its prognosis, and in lung cancer has exhibited disappointing outcomes(24–27). The resistance of PD-1/PD-L1 inhibitors still upset a number of cancer patients. The ability of lung cancer to evade the immune system is main caused by altered cytokines, cellular immune dysfunction, and deficient antigen presentation (28). Due to the lack of predictive biomarkers which limited the application of the immunotherapy in precise medicine of clinical (29, 30), it is urge to find target genes or index that can be used to predicted the response rate of immunotherapy.

Meanwhile, because of the cross-action of tumorigenesis and the immune response, the immune tolerance would develop after receiving immunotherapy(16). In recent decades, next-generation sequencing technology (NGS) has gained popularity and significant success in the advancement of biomedicine (17). In addition, NGS algorithms reveal a wealth of biological information about cancer tissue (18).

The various multi datas clustering algorithm enables to confirmed the news specific molecular subtypes which can rebuilt or find new tissue-based tumor typing. The molecular classification with genetic and epigenetic alterations which can show a new view on drew immune cells infiltration and often provide a predictive value for immunotherapy response. Thus, it is hope to confirm the key genes or index to promote immunotherapy response with based on revealing the connection between the genetic or epigenetic alterations and immune cells infiltration.

Here, our study aim to find immune-related signature genes to establish specific molecular subtypes with different immunotherapy efficiency to get effective and less toxic targeted treatment. We used the multi-omics datas clustering analysis to identified molecular subtypes, and 67 genes was signed as the cluster B signature gene. Based on those genes, we used the ssgsea algorithm to quantify different immune cells infiltration (ICI) subtypes. Our study showed that the score was confirmed as a promising value for predicting immunothrepy efficiency. And meanwhile we used the in-silico drug screening analysis of LUAD cell lines to confirmed the potential drug targets for different ICI subtypes.

## Materials and Methods

### Datasets and Samples

The transcriptome expression data and clinical data were obtained from The Cancer Genome Atlas (TCGA) database (https://portal.gdc.cancer.gov/). The somatic mutation and the clinical pathology characteristic was downloaded from cBioPortal (https://www.cbioportal.org/). A total of 519 LUAD patients were left for subsequent analysis after combining available samples from all omics and clinical features. GSE72094 and GSE68465 were downloaded from GEO datasets and were used to be the external validation cohort.

### Tumor-Infiltrating Immune Cells

We used the 5 published methodologies for decoding tumor microenvironment (TME) contexture:CIBERSORT(4), TIMER(5), xCell(6), MCPcounter(7), EPIC(8).

### Multi omics datas clustering and subtype identification

We used above-mentioned five immune cells infiltration matrix to recognize molecular subtypes. In order to confirmed appropriate the number of molecular subtypes, we evaluated the clustering prediction index(9) and Gaps statistics(10). Meanwhile, based on the five immune cells infiltration matrix, we conducted 10 clustering analysis (iClusterBayes, moCluster, CIMLR, IntNMF, ConsensusClustering, COCA, NEMO, PINSPlus, SNF, and LRA) to confirmed two specially molecule subtypes. Each subtypes silhouette score are similarity.

### WGCNA

The WGCNA package was used to confirmed the related modules about the two clusters(14). The soft threshold was calculated by using the scale-free topology criterion. The optimal soft threshold was selected and the minimum module size was set to 30 genes. Using dynamic tree cut identification module, MEDissThres parameter set to 0.25.

### Differentially expressed genes associated with molecular subtypes

Based on multi omics datas analysis, these patients were divided into two ICI subtypes. The limma package was used to confirmed differentially expressed genes(P<0.01 and absolute fold-change > 1).

### Dimension reduction of Differentially expressed genes and Construction subtypes score

The Boruta package was used to reduce the variables and noise. And the DGEs were positive with the clusters (cor>0.4) was identified as signature A and when it was negative with clusters (cor<-0.4) was confirmed as signature B. Then we used the ssgsea algorithm (11) based on the 67 signature A genes to built grade index to quantify various subtypes.

### Validation set analysis

We used the TIDE website (12) based on the 535 cancer patients from TCGA data to predict immunotherapy responses. Meanwhile we downloaded the Transcriptome data of immune-related genes after PD-1 blockade from IMvigor210 dataset(13).

### Statistical analysis

We used the R software v4.0.3 to conducted the all analyses. Wilcoxon test was be used between two subtypes in continues data. And the pearson correlation coefficient was used to evaluate the relationship between two factors. Kaplan-Meier survival analysis was used to compared different subtypes survival rate. P < 0.05 was considered a statistically significant difference.

## Result

### Identified two molecular subtypes by MOVICS

Based on the five immune cells infiltration matrix, we use multi omics datas clustering to confirmed molecular subtypes with using ten clustering algorithms--iClusterBayes, moCluster, CIMLR, IntNMF, ConsensusClustering, COCA, NEMO, PINSPlus, SNF, and LRA. According to the results of CPI and Gaps analyses, we confirmed the number of subtypes was 2 which the score of the two sutypes was same(figure1A,B,D). The distribution of the various immune cells infiltration matrix is shown in figure1C, and cluster B showed high immune cells infiltration and better survival rate (figure1E). And then the limma package was used to identified the differential expression genes of two clusters(figure1F).

**Figure 1:**
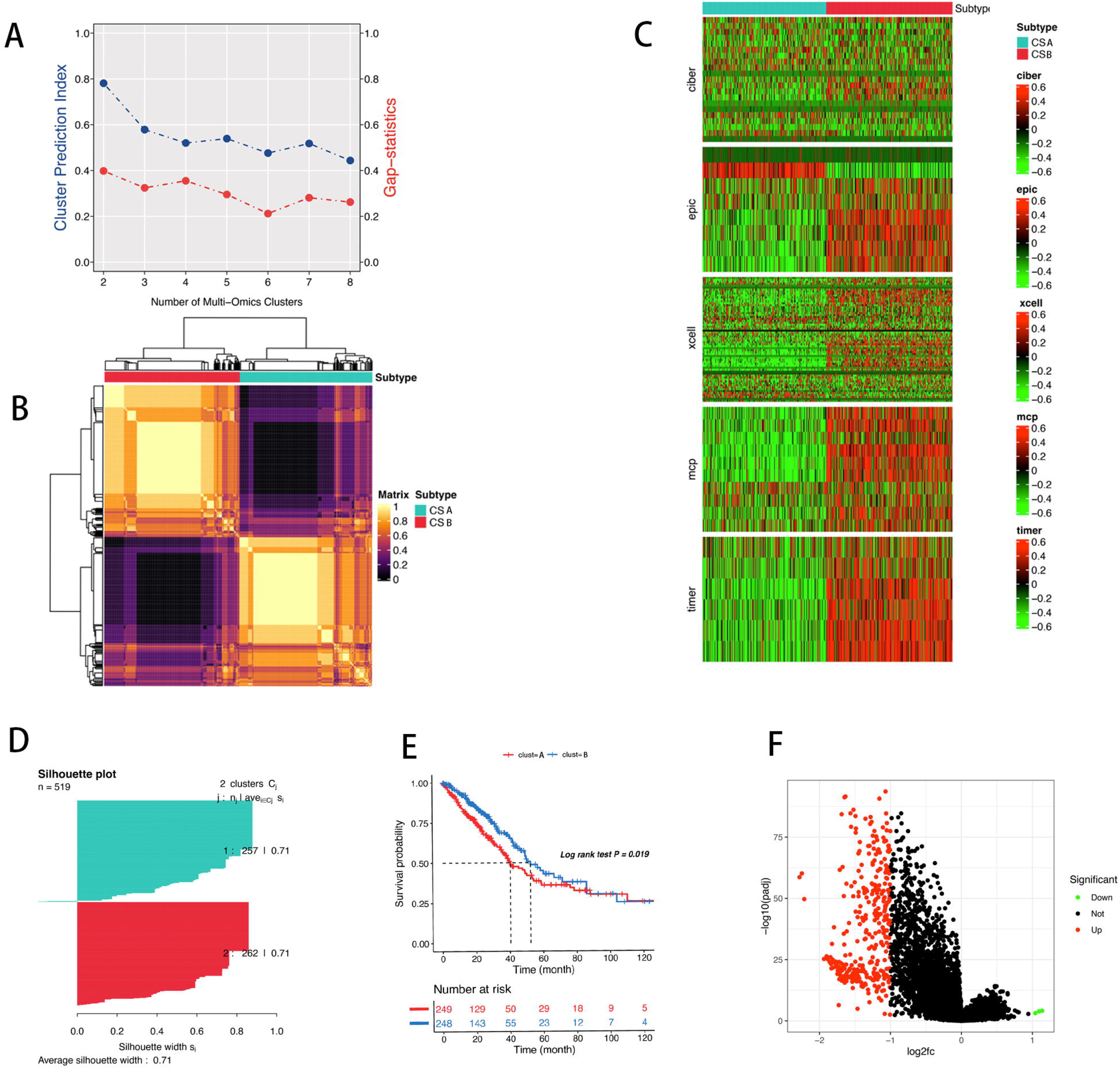
Identifying two molecule subtypes by using various clustering. (A) CPI analysis and gap-statistical analysis results. (B)Consensus matrix based on the various algorithms. (C)The landscape of molecular subtypes in the various immune cells infiltraion. (D) Silhouette-analysis evaluation. (E) The differential survival rate in the two clusters. (F) Differential expression genes in the two clusters.

### Transcript factors and copy numbers analysis

In order to figure out whether the different ICI clusters are the result of different complex regulation molecular network, including genetic and epigenetic event. We evaluated the different activation of 8 LUAD-specific transcription factors and 35 chromatin remodeling potential regulons. And as the figure 2A shown, the cluster A is likely regulated by the EHMT2, HDAC8, KAT2A, ATOH8, SIRT4, KAT5, SIRT5 and FOXA2. And the cluster B differed with higher activity of KDM4C, KAT6B, KDM3B, DACH1, EPAS1, HOXA4, EP300 and KMT2A. The different regulon activity information in the two clusters, which identified epigenetically-driven transcriptional networks were, important differentiators of these clusters. The copy number alteration between the two clusters was assessed, and the result showed that the lost and gained copy numbers were significantly increased in the cluster A compared with the cluster B(figure2B). We observed that the significance lost copy number in cluster A was mostly located at the 9p21.3 region than the cluster B(figure2C,D). IFN family genes play an important role in the regulation of PD-L1 expression (19). High interferon gamma (IFN-γ) levels with accelerated lymphocyte infiltration may be key to identifying tumors with cytotoxic immunophenotypes that are more likely to respond to PD-1 therapy (20). We analysed the quality of CNAs of genes between the two cluster, and the genes located on the 9p21.3 region was significant loss copy number,especially the IFN family genes (IFNB1, IFNW1, IFNA10, IFNA16, IFNA17, IFNA21, IFNA4)(tableS1). Based on that, the cluster A may escape immune surveillance by genomic alterations affecting the copy numebers of the IFN family genes.

**Figure 2:**
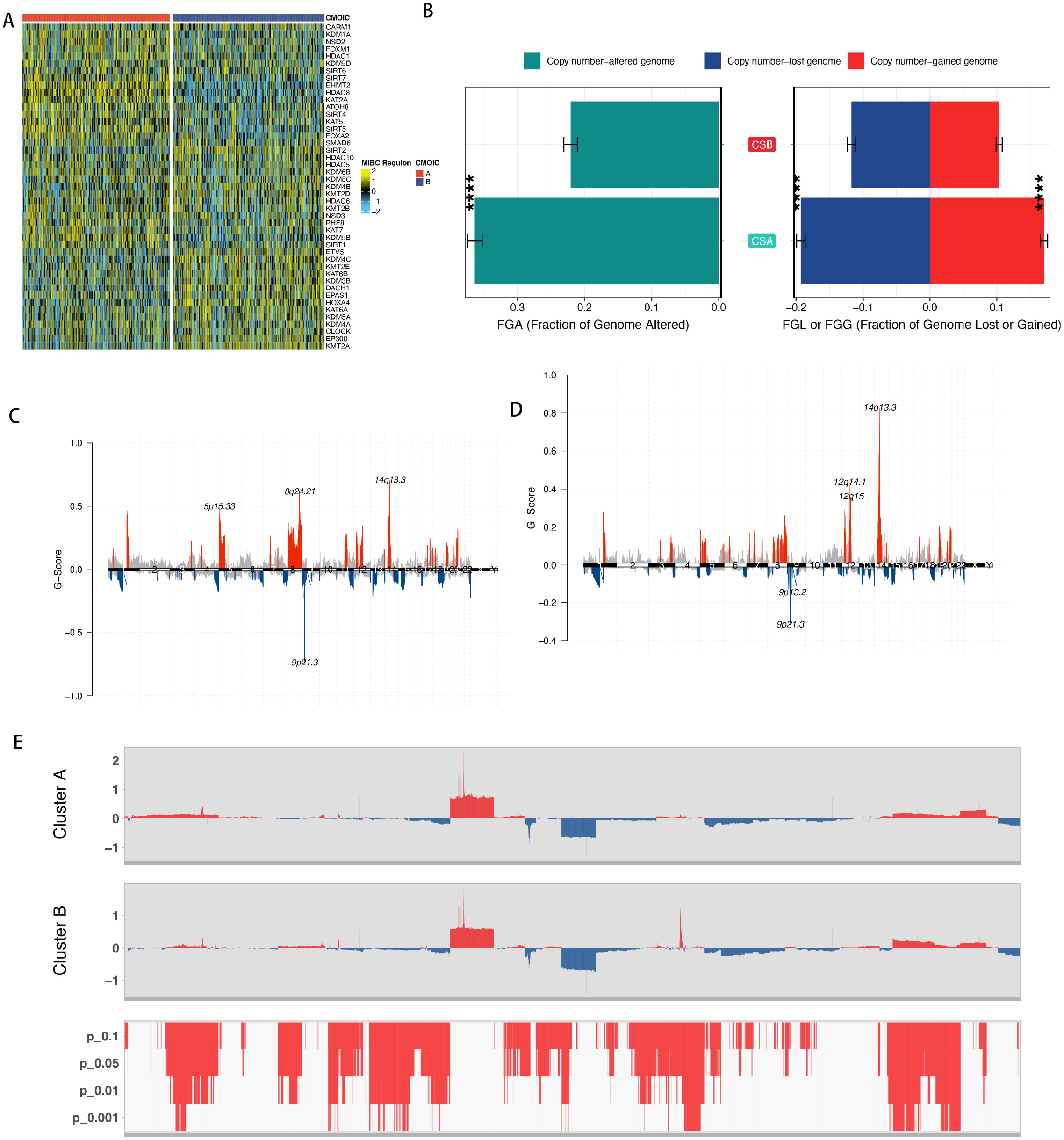
The alternation in the copy number and transcript factors activity. (A) Heatmap showing profiles of regulon activity for 8 LUAD-specific transcription factors and 35 chromatin remodeling potential regulons. (B) The distribution of fraction genome altered (FGA) and fraction genome gain/loss (FGA/FGG) in the to clusters. (C,D) The distribution of cluster A (C) and cluster B(D) in the copy number of genomic regions. (E) CNA plot shows the relative frequency of copy number gains (red) or deletions (blue) between the cluster A and cluster B of the LUAD cohort.

### WGCNA network module mining

Soft threshold values are calculated using the scale-free topology criterion. And as the soft threshold power β = 12(figure3A,B), the phylogenetic trees were then used to mine co-expression modules (cut height≥0.25)(figure3C). By hierarchical clustering analysis of each module, the modules in the same branch showed similar gene expression patterns(figure3D). Then, similar gene modules were identified and combined to obtain 14 co-expression modules (red, blue, green, brown, greenyellow, black, tan, brown, magenta, pink, purple, turquoise, yellow and gray)(figure 3E). Subsequently, the gene clustering was visualized and correlations among the modules were analyzed(figure3F). The magenta module has high correlation with the cluster B. The module is selected as an important module for further analysis (Figure 3G)(tableS2).

**Figure 3:**
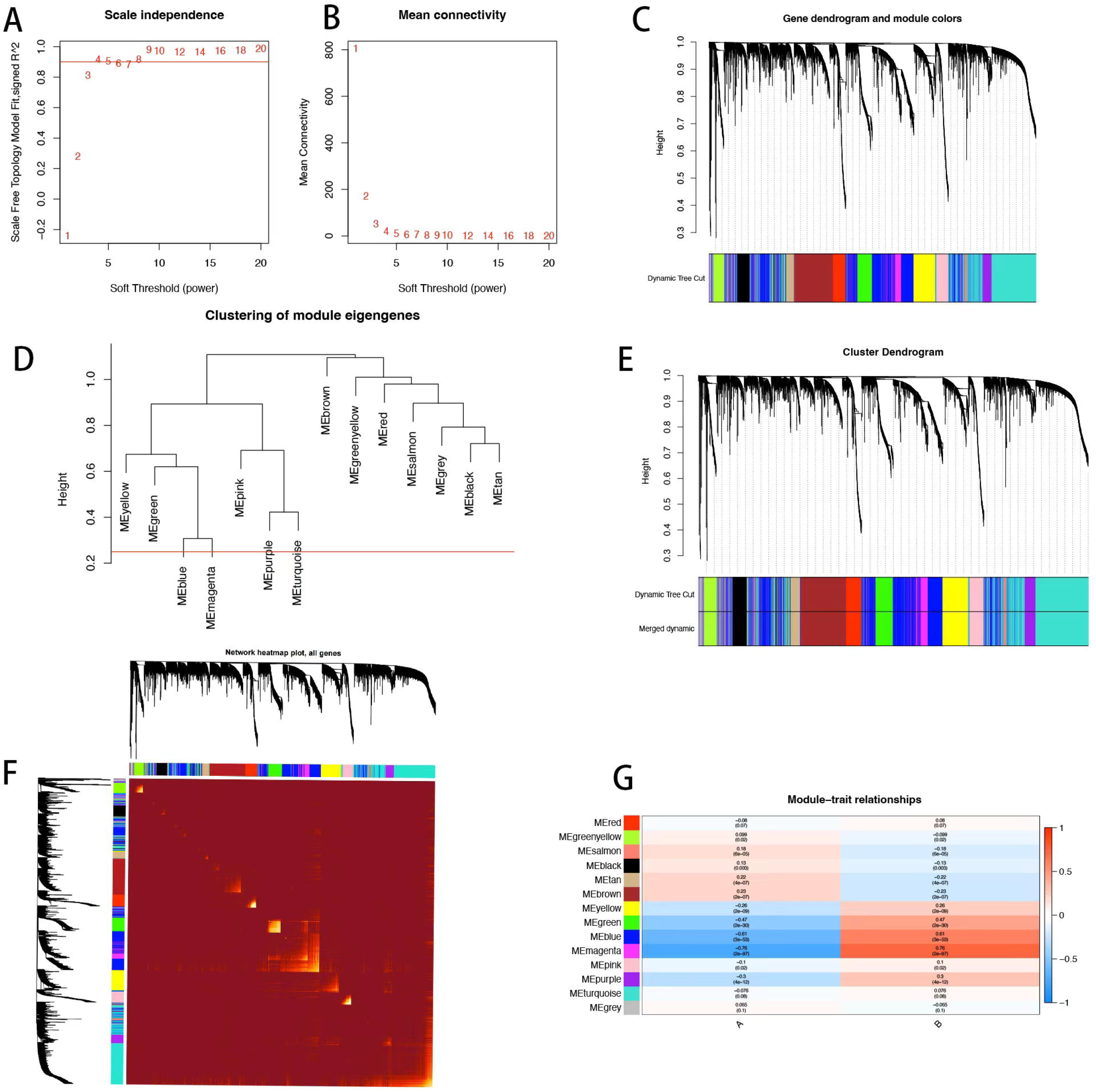
WGCNA analysis for the two clusters. (A) Confirming the best scale-free index for various soft-threshold powers (β). (B) The mean connectivity for various soft-threshold powers. (C) The gene tree map and nodule color. (D) Hierarchical clustering analysis. (E) The gene dendrogram is based on clustering. (F) The heatmap of all genes. (G)Heatmap of the correlation between the module genes and the two clusters.

### Dimension reduction and Construction of the signature genes Score

In order to evaluate the different biology function between the two clusters, the differential activation level of 50 tumorigenesis-associated pathways in the two clusters was analysed. And the result showed that the significance activation of the G2M checkpoint, E2F target pathways and DNA repair in the cluster A, and the cluster B had high activation level in the IL6 JAK STAT3 signaling, interferon alpha response and interferon gamma response(Figure 4A). The above result suggested the high activated of cell proliferation-associated pathways were correlation with poor prognosis. And based on those DEGs and the magenta module genes, the pearson correlation coefficient was used to evaluate the relationship between genes and clusters. And then we used boruta package to reduce the dimension in both signature A and B for reducing the noises of cluster genes. We confirmed 67 genes as signature A gene(cor>0.4,p<0.01), and signature B genes were not meet those conditions(figure4B)(tableS3). As the figure4F shown that the signature A genes were enriched in the positive regulation of T cell activation, leukocyte migration and regulation of T cell activation. Then we analysed the correlation between the clinicopathologic factors and the two clusters. The chi-squared test showed that the cluster A exhibited poorer TMN staging and survival status than the cluster A, suggesting that the ICI clusters had high predicted values which is associated with highly malignant LUAD(figure4D). These results indicated that our model is a good indicator for prognosis and survival in LUAD. Based on the signature A genes, we used the ssgsea algorithm to count the score which can quantify the subtypes. As the figure4B shown that the high score is overlap with most of cluster A. And then we evaluated the different score groups survival rate and the result showed the rate of high score group is higher than the low score group(figure4D). We selected 13 immune related genes (CD274、CD8A、CXCL10、CXCL9、GZMA、CTLA4、 HAVCR2 、 IDO1 、 LAG3 、 PDCD1 、 PRF1 、 TBX2 and TNF) as the immune-activity-related signatures and the high score group showed high immune activity(figure4F).

**Figure 4:**
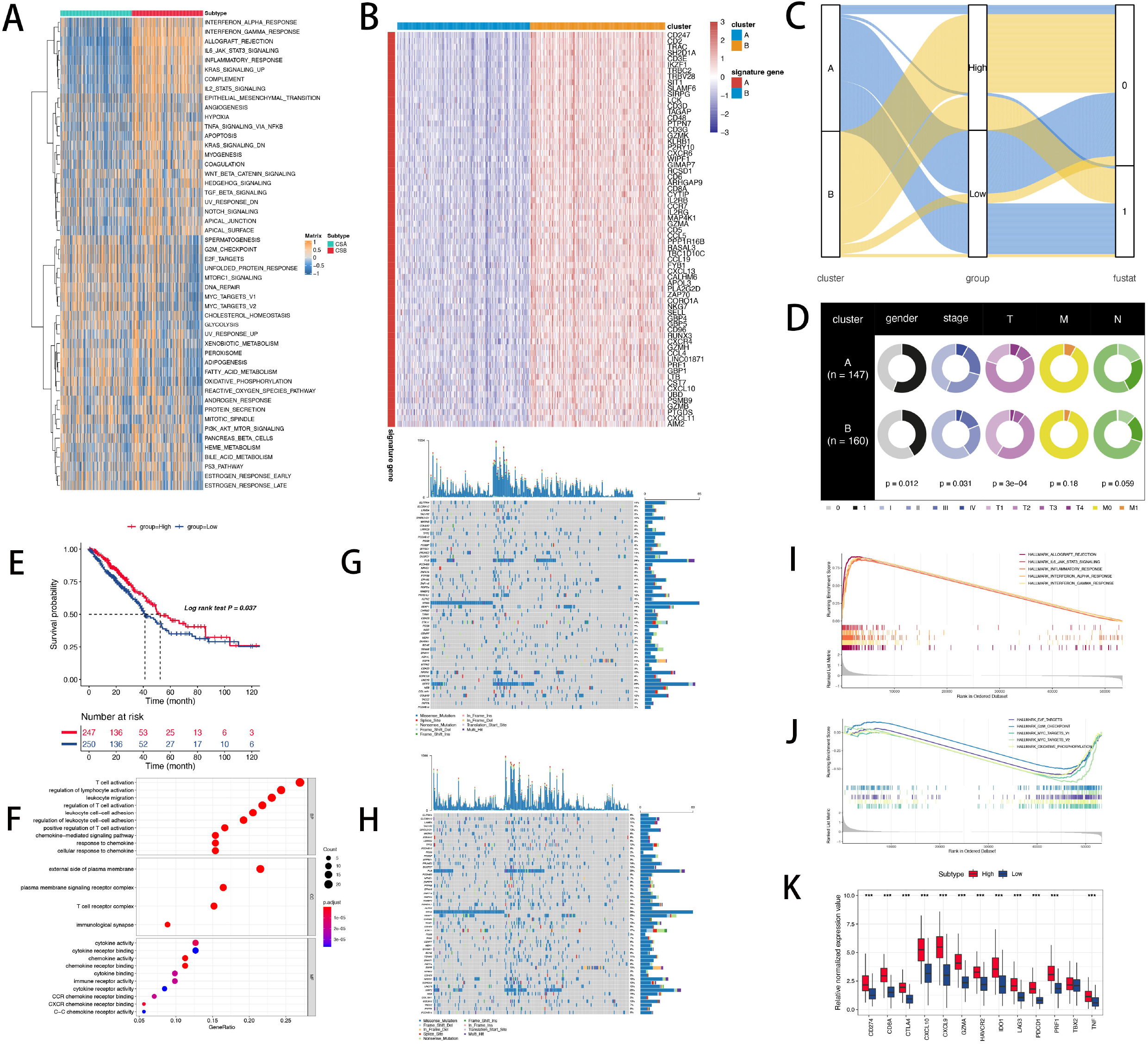
The alternation of genetic and mutation in the different subtypes. (A) Heatmap of 50 HALLMARKER pathways in the two clusters. (B) The clustering analysis in DEGs and magenta module gene of two clusters. (C)The distribution of clusters, and survival outcome in different score groups. (D) The distribution of clinicopathologic factors in the two clusters. (E)The differential survival rate in the different score groups. (F)The biological function of the signature A genes. (G,H) The significant mutation genes in the high scores (G) and low scores(H). (I,J)The biological function of low and high score groups. (K)The distribution of immune-related genes expressed in high and low score groups.

### The distribution of immune cells infiltration of subtypes in external cohorts

In order to get the score in the external cohort (GSE68465 and GSE72094),we used the 67 genes(signature A genes) to conducted the ssgsea algorithm. In the two cohort, the high score group is the inflammatory subtype and the low score group can be confirmed as desert immune cluster(figure5A,B). Those results are consistent with the TCGA cohort(figure1B).

**Figure 5:**
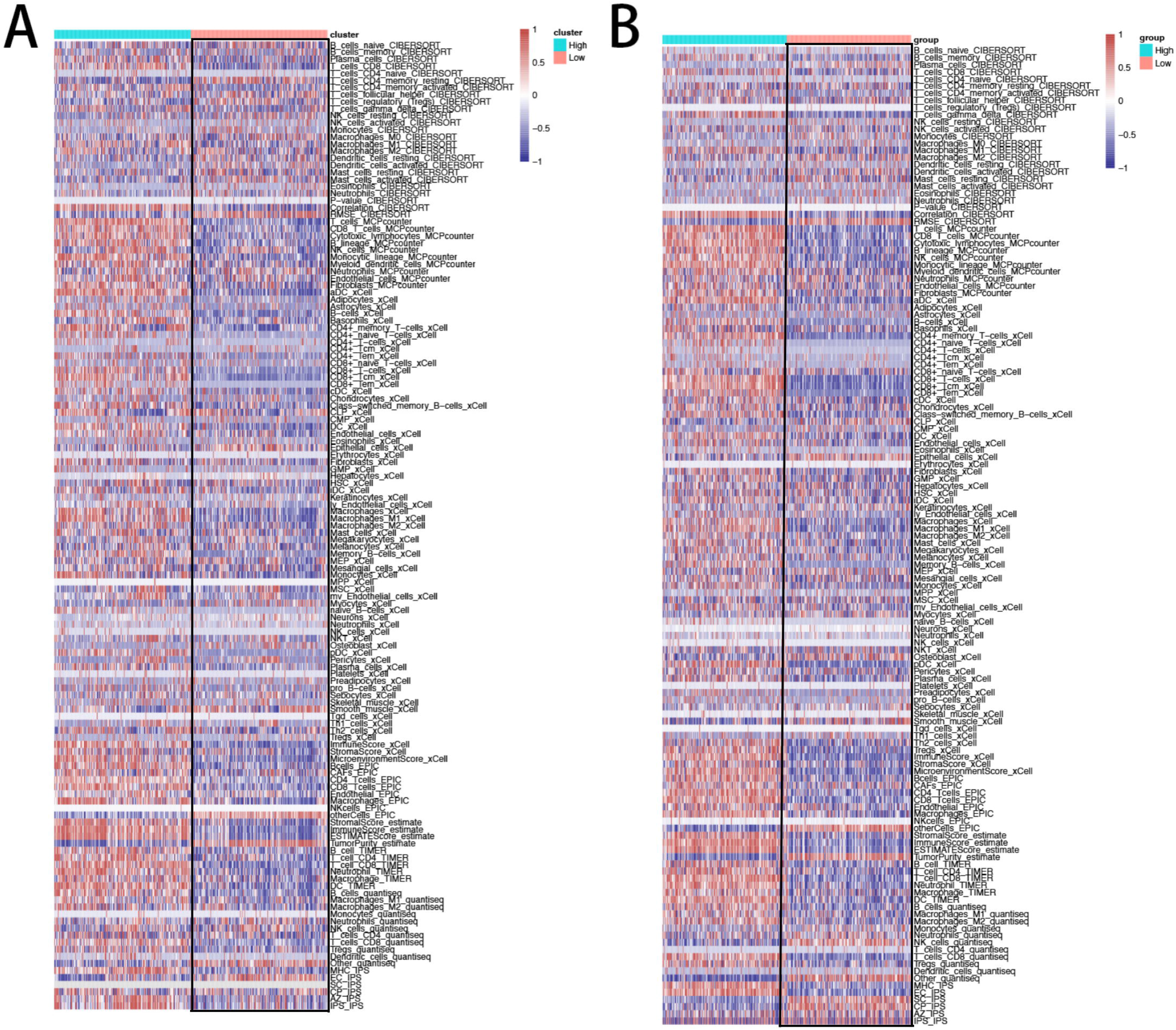
Validation of different score group in external cohorts. (A,B) The distribution of various immune cells in the GSE72094 data (A) and GSE69465 data (B).

### The value of the Scores in the Prediction of Immunotherapeutic Response

The applying of immunotherapy had greatly improve the cancer patients survival rate, especially the use of PD-1 or PD-L1 specific monoclonal antibodies. While, the remission rates of immunotherapy remain lower than expected, rarely exceeding 40 percent. Here, we analyzed the value of the score in the immunotherapy with IMvigor210 cohort and TCGA cohort. Based on the 67 signature A genes, we used the two cohort conducted ssgsea algorithm and evaluated the effect of anti-PD-L1 immunotherapy of two score groups. The TIDE website (12) was used to predict the cancer immunotherapy responses of 535 patients in the TCGA database. From the perspective of prediction results, the objective effective rate of anti-PD-L1 treatment of TCGA in high score group (21.1%) was higher than that in low score group (0.4%) (Figure 6B) and the IMvigor210 cohort had the same trend with the TCGA cohort in the objective effective of anti-PD-L1 treatment(high score:28.8%, low score:18.8%)(figure6E). The subclass mapping was used to compare the expression profiles of the two score subtypes and the result showed the score group treatment respond to anti – PD-1 therapy (Bonferroni correction P<0.01) (figure6C,F). Meanwhile the response subtype had high score than the non-response subtype (Figure 6A,D).

**Figure 6:**
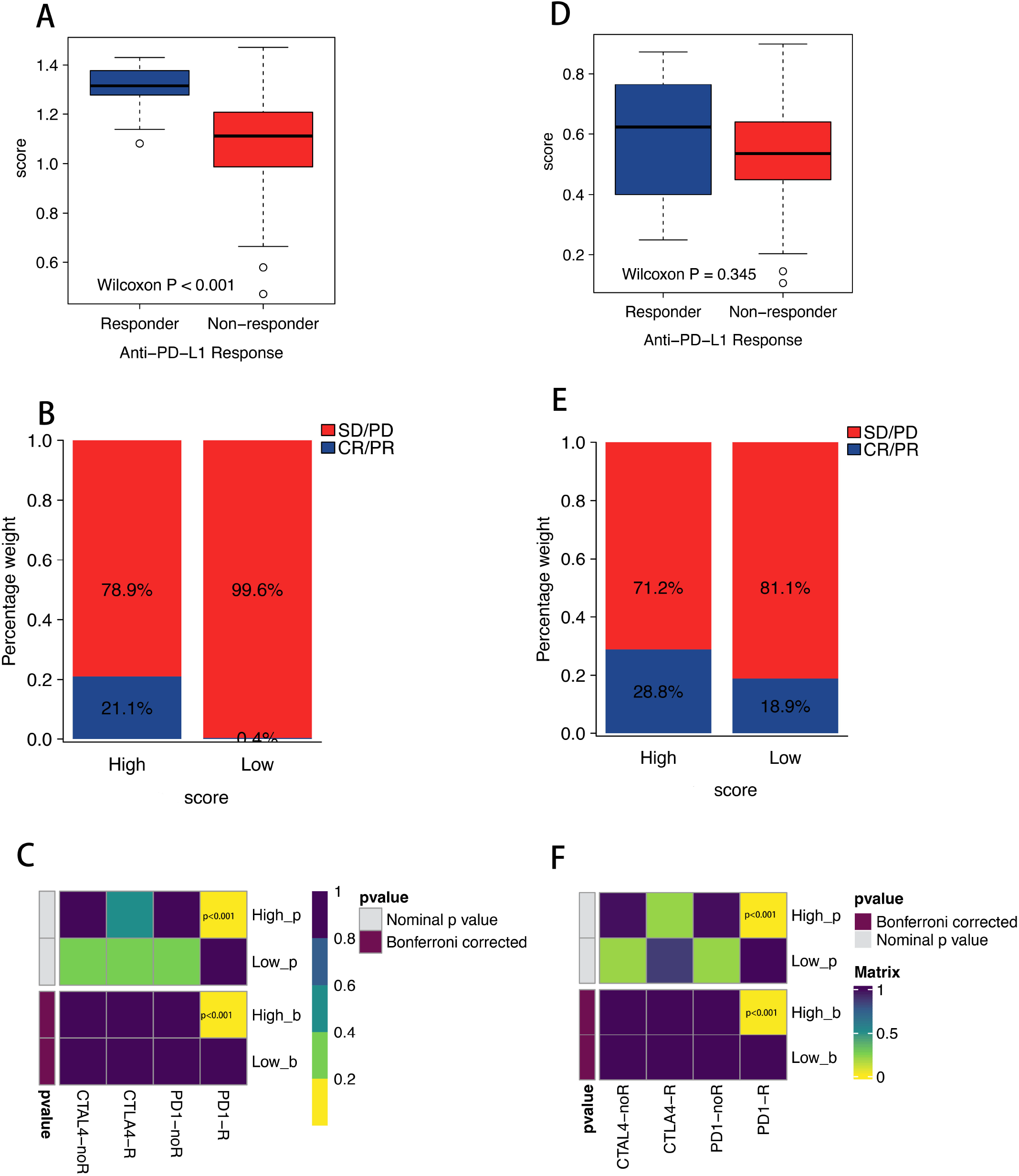
Differential response to immunotharepy in the different score groups. (A,D)The distribution of score in the different anti-PD-1 response. (B,D) The response proportion of anti-PD-L1 immunotherapy (response/Non-response and stable disease (SD)/progressive disease (PD)) in the different score groups in the Imvigor210 and TCGA cohort. (C,F) Differential immunotherapeutic response in the high score group and low score group.

### In-Silico Drug Screen for lung cancer cell lines Resembling a High or Low score Profile

Then we used the GDSC data to select the compounds which are sensitivity to the different score subtypes to elucidate for a more effective treatment of high score group and low score group. Based on the 67 signature A genes, we conducted the ssgsea algorithm with the lung cancer cells lines to built score. And interesting, the 67 signature A genes exhibited a significant expression in the transcript levels in lung cancer cells lines(figure7A). The drug sensitivity analysis showed that the high score group revealed a significantly higher sensitivity of LGK-974 and the low score group can benefit more from the EGFR target therapy (erlotinib) (figure7C)(tableS4).

**Figure 7:**
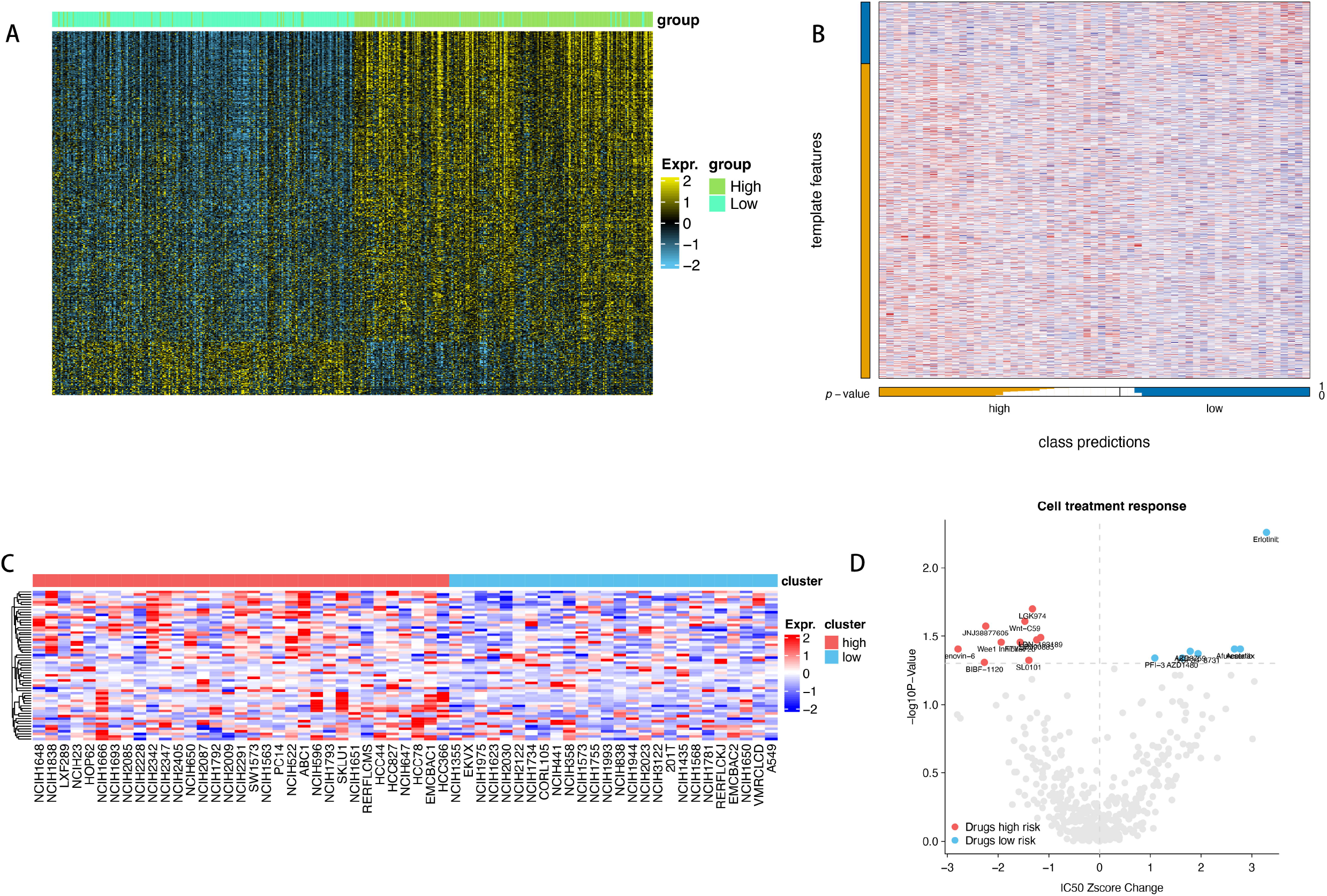
EGFR inhibitors as potential drugs for low score group. (A)Heatmap of subtype-specific upregulated biomarkers using limma for identifying the two clusters. (B) Heatmap of NTP. (C) Heatmap shows the expression pattern of 67 genes in cancer cell lines showing low or high score group. (D)The figure summarizes the relative changes of the ic50z score and P values in selected luad cell systems at low or high score group which can be treated using the designated compounds based on the cancer drug susceptibility genomics (gdsc) database. The red spot clearly shows the high sensitivity drugs of high score group and the blue point clearly shows the high sensitivity drugs of low score group.

## Discussion

As the World Health Organization research showed that the cancer is the second leading cause of death worldwide, and the lung cancer is the main reason for causing death in the cancer-related death. The Non-small cell lung cancer is the main subtype of lung cancer including squamous cell carcinoma, adenocarcinoma and large cell carcinoma. Recently, the immunotherapy had been widely used in clinical practice to improve the prognosis of various cancers. However, the effective response rate of PD-1 inhibitors is still low. Previous studies mainly focused on some specific genes expression for predicting the prognosis of tumors (31,32,33). However, the biological process of tumors is extremely complex, and the characteristics of different types are connection with each other. Using multi-omics analysis to explain the tumors heterogeneity and construct relevant models to predict the efficacy of immunotherapy can make the system more stable and convincing. This process is not only reduces the inauthenticity of CIBERSORT’s results (34), but also can make the model system stable and not be influenced by the expression of single or multiple genes (35,36,37). We try use ten clustering algorithm to identified two clusters, and the survival outcomes of the two ICI subtypes were significantly separated, as were the specific molecular features. First, we used five method to construct the the landscape of immune cell infiltration in LUAD, and we identified the two different immunological status of tumor with the clustering. The cluster B showed a favorable prognosis, as well as higher infiltration percentage of most immunological cells. The previous research has been demonstrated the different ICI subtypes correlated with the cancer patients survival rate (39). According this finding, our study showed that the cluster B had a favorable prognosis of LUAD patients which may attribute to the higher immune cells infiltration which can be identified as “hot” immunological status may be more easily for immune therapy (38) consistent with the previous study. The cluster A exhibited the worst prognosis which may be influence by the activated cell proliferation pathways and regulated by the EHMT2, HDAC8, KAT2A, ATOH8, SIRT4, KAT5, SIRT5 and FOXA2. Also the cluster A had instability genetic alteration, especially the copy number amplification of 9p21.3. For the amplification, we observed that the IFN family genes located in the 9p21.3 was significant loss in the cluster A than the cluster B. As the previous reported that copy number loss in IFN-γ pathway genes can work as a predictive factors fro predicting the immunotherapy response(40,41,42). Combined with our results, the loss of IFN-γpathway genes may be the main reason of cluster B to evade immune surveillance.

Then we used the WGCNA analysis and limma package to fetched the module genes and differential expression genes between the two clusters and the boruta package to reduce the dimension in both signature A and B. Based on signature A genes which was enriched in the T activated pathways, we used the sssgsea to quantify the ICI clusters. We divided those patients to the high and low score based on the score median value. And the high score is most overlap with the cluster B, which consistent with our thoughts. The high score group exhibited better survival rate and the high activated of IL6 JAK STAT3 signaling, interferon alpha response and interferon gamma response. In our study showed that the high score had high expression of immune-activity-related signatures which had huge potential benefit and response from immunotherapy. We used the two external cohort (GSE68465 and GSE72094) to verify our mode accuracy in predicted the immune cells infiltration distribution in the high and low score group. Lung cancer is the most commonly cancer worldwide. Nowadays, surgery is still the primary treatment for lung cancer, however when it is diagnosed as the advanced malignant tumor, the chemotherapy and radiotherapy are the next options to be considered in such cases, these forms of treatment have side effects that can sometimes be fatal to patients. Therefore, new and effective strategies with minimal side effects are urgently needed. Cancer immunotherapy is now a promising to cure cancer or improve the prognosis of cancers. Multiple immunotherapy agents have been proposed and tested for potential therapeutic benefits in lung cancer, and some have fewer side effects than conventional chemotherapy and radiation therapy. In our study, the high score group can be identified as the most promising immunotherapy subtypes. And we used the IMvigor210 cohort and TCGA cohort to conducted the correlation with the score. The results showed that in the two cohorts, the high score had rate of anti-PD-L1 therapy than the low group, as well the submap result showed that the high group have a high likelihood of responding to immunotherapy.

Also, the drug screening predicts that the elefinib, an EGFR inhibitor, is an effective compound for low scores group. Many studies have reported the association between EGFR MEK pathway activity and the immune cold phenotype of multiple cancers, including LUAD, and the advantage of ICI treatment in EGFR mutation is low (21). Our result showed that the low score group might benefit from the targeted inhibition of the EGFR.

In conclusion, we built a new novel signature score for LUAD patients via multi-omics data and consensus results of ten algorithms. The high score group identified as the activation of inflammation-related pathways and high expression of immune checkpoint proteins with better survival. Meanwhile the high score group have a high likelihood of responding to immunotherapy. Our model can be effective to predicted the efficacy of anti-PD-L1 and prognosis. However the score model should be confirmed by a prospective analysis of a large LUAD patient cohort, and the efficacy of anti-PD-L1 for high score requires validation in appropriate pre-clinical models and future clinical trials.

Our data emphasized the complex interactions between genetic and apparent genetic events in the construction of tumor immune phenotypes, suggesting that the LUAD patients with high scores failed in ICI treatment may benefit from the combination of EGFR inhibition.

## Notes

### Competing Interest Statement

The authors have declared no competing interest.

